# The Physical Basis of Osmosis

**DOI:** 10.1101/2023.01.02.522450

**Authors:** Gerald S. Manning, Alan R. Kay

## Abstract

Osmosis is an important force in all living organisms, yet the molecular basis of osmosis is widely misunderstood as arising from differences in water concentration in solutions of differing osmolarities. In 1923 Debye proposed a physical model for a semipermeable membrane that was hardly noticed at the time and slipped out of view. We show that Debye’s analysis of van’t Hoff’s law for osmotic equilibrium provides a consistent and plausible explanation for osmotic flow. A difference in osmolyte concentrations in solutions separated by a semipermeable membrane generates different pressures at the two water-membrane interfaces. Water is therefore driven through the membrane for exactly the same reason that pure water flows in response to an imposed hydrostatic pressure difference. In this paper we present the Debye model in both equilibrium and flow conditions. We point out its applicability regardless of the nature of the membrane with examples ranging from predominantly convective flow of water through synthetic membranes to purely diffusive flow of independent water molecules through a lipid bilayer and the flow of strongly interacting water molecules in single file across narrow protein channels.

## Introduction

Osmosis is one of the most powerful forces that organisms must counteract to survive. An index of its importance is that animal cells, of all kinds, spend about a quarter of their energy resisting the osmotic challenge induced by the presence of impermeant molecules in cells (*i*.*e*. the Donnan effect, Appendix 1) (Rolfe and Brown, 1997; Kay, 2017). An unchecked Donnan effect would lead to the continuous influx of water until the cell bursts. The need to maintain osmotic balance is unrelenting, interrupted neither by sleep nor hibernation. Furthermore, osmosis is quite literally at the root of all plant function (Niklas and Spatz, 2012). Yet, although the phenomenological thermodynamics of osmosis has long been clear, at least for osmotic equilibrium, its molecular basis continues to be widely mischaracterized and hence misunderstood, although a consistent mechanistic understanding was presented a hundred years ago (Debye, 1923b).

In this paper we will show why a molecular basis for osmosis that is most often given in biology textbooks is invalid. We will then show how a physical mechanism that was first presented by Peter JW Debye in 1923 (Debye, 1923b) can generate a macroscopic pressure and provides the most plausible account of osmosis. We refer to it as the *Debye model*. Debye was perhaps the first to recognize that osmosis arises from the mechanical interaction of an impermeant solute with a semipermeable membrane, but does not depend on the precise chemical nature of the solute or the solvent. We believe that the Debye model has failed to take hold in biology for several reasons, *inter alia*, a lack of understanding of the physical argument, its requirement for mathematical explication, and the availability of other simple, seemingly plausible, but flawed arguments. In addition, textbooks, besides omitting the Debye model, have not raised any inconsistencies in the conventional approach. There has hence seemed little need to question what at first blush seems a simple phenomenon.

There have been several attempts primarily directed at biologists to set the record straight on the physical basis of osmosis (Stein, 1966; Kiil, 1982; Kramer and Myers, 2012), as well as accounts of the Debye model in journals (Manning, 1968; Oster and Peskin, 1992; Borg, 2003; Marbach and Bocquet, 2019; Song et al., 2021) and textbooks (Benedek and Villars, 1974; Weiss, 1996; Baumgarten and Feher, 2011; Nelson, 2014), but despite these efforts, misconceptions have persisted. The apparent simplicity of osmosis may have masked what is at bottom a rather subtle phenomenon with enormous implications for biology (Dick, 1966; House, 1974; Andersen, 2015). It is we think worth readdressing the physical basis of osmosis because it may open new ways of looking at water and solute transport that have remained hidden because of flawed beliefs.

The osmotic flux of water is important in several biological disciplines; indeed it is a challenge to find one where it is not. However, different branches of science have developed unique terminologies, which may confuse someone familiar with the terms of one field in reading the literature of another. The unified view and terminology presented here may help to bring consilience to the study of osmosis.

We first provide a review of the basic empirical knowledge of osmosis, including a discussion of some misconceptions. Then we give an account of the Debye model, both as presented by Debye himself to derive van’t Hoff’s law for osmotic equilibrium, and as extended to apply to osmotic flow (Manning, 1968). Finally we use Debye’s model to deepen our understanding of water flow across biological membranes, specifically, diffusion across lipid bilayers and in single file across nanopores such as aquaporin.

### 1 The rudiments of osmosis & common misconceptions

**Table 1:**
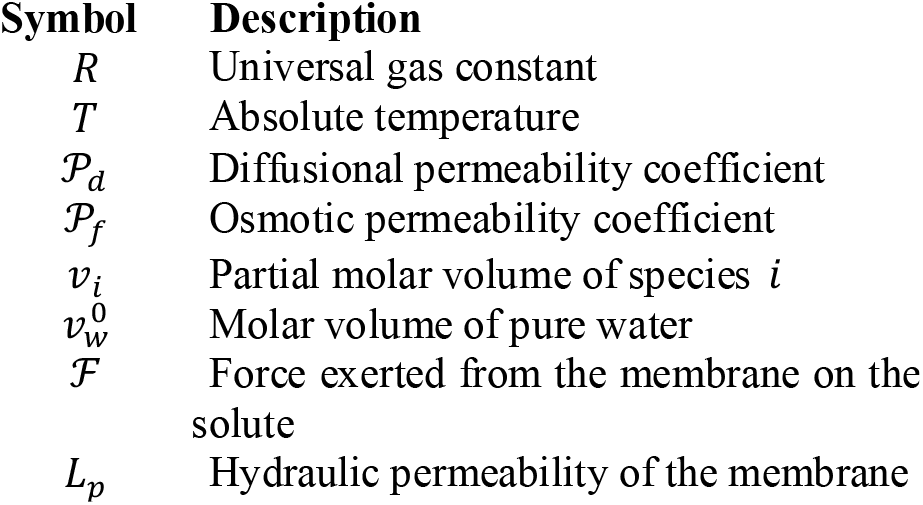
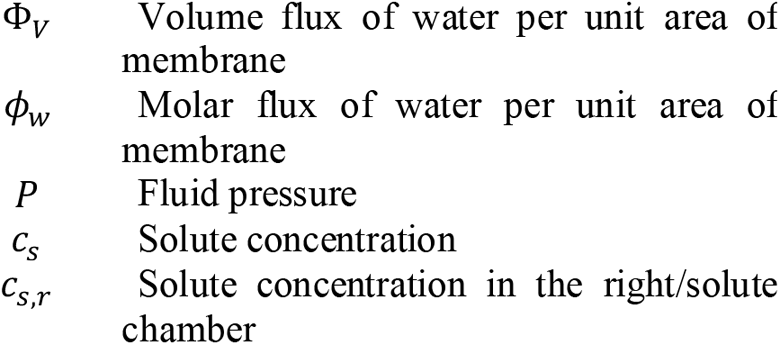
Symbols used in the text

To illustrate the process of osmosis we will consider a **semipermeable membrane**, namely, one that is permeable to water but completely impermeable to solute molecules, separating two solutions. We will restrict our discussion to water, but it also applies to any other solvent. If the **osmolarities** (*i*.*e*., the total concentrations of solutes) of the solutions differ, water flows from the solution with the lower osmolarity to that with the higher osmolarity. In the situation diagrammed in Fig. 1, the movement of water can be prevented if the piston exerts an excess pressure on the solution with higher osmolarity equal, if the solutions are dilute, to *RT*Δ*c*_*s*_, where Δ*c*_*s*_ is the osmolarity difference. This experimental observation is encapsulated by van’t Hoff’s equation,

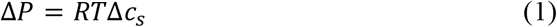

where Δ*P* is the pressure difference in no-flow, equilibrium, conditions between two solution chambers separated by a semi-permeable membrane. ^1^

**Figure 1:**
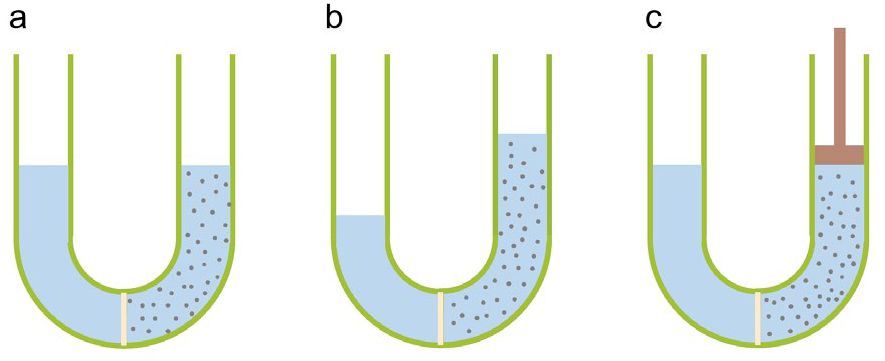
Classical demonstration of osmosis. (a) A U-tube with a semi-permeable membrane separating pure water on the left from an aqueous solution with an impermeant solute of concentration Δ*c*_*s*_ on the right. (b) With time water will move from left to right, elevating the column of solution on the right, until the gravitational weight imbalance stops the flow. (c) Alternatively the flow of water can be prevented if a piston applies a pressure equal to *RT*Δ*c*_*s*_ (in the dilute regime).

The term *RTc*_*s*_ in a free-standing solution with solute concentration *c*_*s*_ is often referred to as the “osmotic pressure” of the solution. However, this imprecision is the source of much confusion since an actual osmotic pressure difference can only arise between two solutions with different osmolarities separated by a semipermeable membrane. *It is worth emphasizing that osmotic pressure is not a physical property of a free-standing aqueous solution*.

Our objective is now to understand *what* generates such a pressure difference across a semipermeable membrane separating solutions with different osmolarities. Van ‘t Hoff’s law can be derived by equating the chemical potentials of the water in the two compartments (Dill and Bromberg, 2003; Phillips et al., 2012), but the thermodynamic derivation provides no insight into the molecular mechanisms that generate the pressure difference. To begin our analysis, we review first the hydraulic flow of water in response to a hydrostatic pressure difference, and relate this motion to that induced by a difference in osmolarity. We consider a membrane with pure water on both sides when a transmembrane **hydrostatic pressure difference** Δ*P* is imposed (for example with a piston). The **volume water flux per unit area of membrane** is given by the empirical relationship(Weiss, 1996).

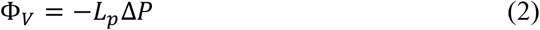

with the water flux being directed to the side with lower pressure, and *L*_*p*_ is called the hydraulic permeability. The value of *L*_*p*_ depends on the specific composition and structure of the membrane that allows water to move across it. (2) is Darcy’s law, which can be derived from the Navier-Stokes equation for the convective flow of a liquid.^2^

The volume water flux across a semi-permeable membrane subject to both a hydrostatic pressure difference and a difference in osmolarity can be derived by combining Eqs. (1-2),

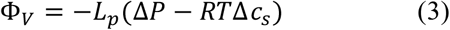

(3) has a long history and has been proposed by many scientists in different fields, sometimes only in words. It is often called Starling’s equation in physiology (Starling, 1896; Blaustein et al., 2019) and it is part of the Kedem-Katchalsky (1958) equations, but it could without exaggeration be called the **Fundamental Law of Osmosis**.

A remarkable feature of Eq. 3 is that two physically distinct driving forces, an imposed hydrostatic pressure difference Δ*P* and an osmolarity difference *RT*Δ*c*_*s*_, produce the same flux of water. The connection between force and flow is given by the same coefficient *L*_*p*_ in both cases. The implication for the underlying physical mechanisms of pressure and osmotic flow is that these mechanisms must be one and the same.

Note also that van ‘t Hoff’s law at equilibrium is recovered from the Fundamental Law by setting the flux Φ_*V*_ equal to zero. If the coefficients for the two driving forces were different, van ‘t Hoff’s law would be violated.

When the volume flux is carried only by the water, the number of moles of water flowing across unit area of membrane can be derived for dilute solutions from the molar volume of water 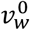 (Finkelstein, 1987).

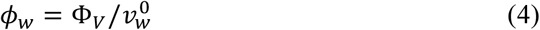

Substituting equation 3 into 4, gives an alternative form of the Fundamental Law of Osmosis,

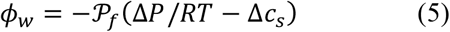

where 𝒫_*f*_ is the osmotic permeability coefficient,

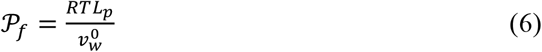

𝒫_*f*_ can be determined from the measurement of water fluxes induced either by a hydrostatic pressure difference or a difference in osmolarity across a membrane (Fettiplace and Haydon, 1980; Finkelstein, 1987).

The foregoing observations give rise to several questions, which we will pick up later. What is the physical reason for the observed equivalence of hydraulic and osmotic flow? It is counter-intuitive that the same coefficient *L*_*p*_, or 𝒫_*f*_, should apply to both. Why, at equilibrium, must an impermeable solute concentration be balanced by a difference in hydrostatic pressure, and why should van’t Hoff’s law be so similar to the equation of state of an ideal gas?

#### 1.2 How water moves across membranes

The flow of water is composed of two components, a **convective** component and a **diffusive** component (Truskey et al., 2009). Both may be present simultaneously but to different degrees depending on the nature of the flow. For macroscopic flow, the convective movement dominates, but we will give an example of flow through a lipid bilayer that is entirely diffusive. We describe the convective and diffusive contributions in turn.

**Convection** is the bulk flow of liquid induced by a force. It is what we are able to see when water runs in a brook or through a pipe and is described mathematically by the Navier-Stokes equation) (Truskey et al., 2009; Phillips et al., 2012). At the molecular level, in convective flow, clusters of closely packed water molecules move in concert in the direction of the force. However, because molecules in a liquid can move relative to each other, they are always in random motion, which drives diffusive movement. If, in addition to thermal motion, a mechanical force *ℱ* acts on the molecules, their random movements are biased in the direction of the force, and each molecule acquires a **drift velocity** *µℱ* in the direction of the force. The proportionality constant *µ* is called the **diffusional mobility** of the molecule,^3^ and it is connected to the diffusion constant *D* through the Einstein relation *µ* = *D*/*RT*. Molecules within a liquid flowing convectively under a force therefore simultaneously exhibit diffusive motion that is superimposed upon the convective flow. More explicitly, the average velocity of a molecule in a flowing liquid is the sum of the convective velocity and the diffusive drift velocity.

As a pertinent example, a pressure gradient in water simultaneously induces both convective flow according to the Navier-Stokes equation and a diffusive drift of water molecules along the gradient. Clusters of water molecules move as a whole along the pressure gradient, while each individual molecule responds to the gradient by drifting stochastically away from regions of higher pressure and towards regions of lower pressure. The reason that an individual molecule drifts towards a region of lower pressure is that less work is required at lower pressure to accommodate the molecular volume.

For flow through membranes we can quantify the relative importance of the convective and diffusive contributions with the dimensionless 𝒫_*f*_/𝒫_*d*_ ratio. The overall permeability 𝒫_*f*_ has already been defined as characterizing the flow observed when a pressure or osmolarity difference is imposed on the two sides of the membrane in accordance with the Fundamental Law of Osmosis, Eq. 5. The quantity 𝒫_*d*_ is what the permeability would be in the absence of convection. Then, only the diffusivity of the water molecules is effective in the flow. Significantly, 𝒫_*d*_ can actually be measured in a separate experiment from the observed diffusional flux 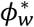 of trace concentrations of isotopically labeled water, (Mauro, 1957; Fettiplace and Haydon, 1980) in the absence of either a pressure or osmolarity difference,

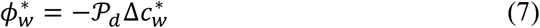

and 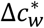 is the difference in concentration of the water isotope across the membrane.^4^ That the 𝒫_*d*_ in this equation is the same 𝒫_*d*_ appearing in the 𝒫_*f*_ /𝒫_*d*_ ratio requires proof, provided in Appendix 2.

It is likely that the diffusional and convective flows of water are additive, so we write 𝒫_*f*_ = 𝒫_*c*_ + 𝒫_*d*_, where 𝒫_*c*_ is the contribution from convection, then divide both sides by 𝒫_*d*_, we find that for the 𝒫_*f*_ /𝒫_*d*_ ratio,

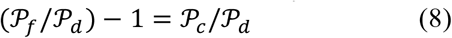

from which a useful interpretation of 𝒫_*f*_ /𝒫_*d*_ emerges. Since from its meaning, convection is represented by a positive value of 𝒫_*c*_, the smallest possible value of 𝒫_*f*_ /𝒫_*d*_ is unity, and then the flow is entirely determined by the diffusivity of the water molecules. But if 𝒫_*f*_ /𝒫_*d*_ is much greater than unity, convection dominates osmotic flow through the membrane.

Mauro (1957) and Robbins and Mauro (1960) measured 𝒫_*f*_ and 𝒫 for a series of synthetic collodion membranes of increasing density in polymer material. For the most open membrane, the diffusive component of water flow was a small fraction, 1/730, of the overall observed flow, while for the most dense membrane the diffusive contribution was somewhat more important, but still just 1/36 of the total. Their experiments showed conclusively that the water flow in these membranes is dominated by convection, like water running in a brook, perhaps obstructed in its course by rocks (in the membrane, by polymer material).

Unlike most synthetic membranes, biological membranes are heterogeneous, with protein channels like aquaporin spanning the lipid bilayer (White et al., 2022). Water is transported independently through both the bilayer and the channels, as illustrated in Figure 2. The 𝒫_*f*_ /𝒫_*d*_ ratio provides insight in the biological case also. For isolated lipid bilayers, measurements show 𝒫_*f*_ /𝒫_*d*_ = 1, so there is no convective flow component. Water crosses the lipid bilayer diffusively, as widely dispersed independent molecules. However, the measurements of Hevesy et al. (1935) in frog skin many years ago showed that 𝒫_*f*_ was greater than 𝒫_*d*_. This inequality was also found to be true in red blood cells (Paganelli and Solomon, 1957). These experiments provided the first evidence of water channels; however, it took a long time to identify and isolate aquaporin channels (Agre et al., 1995).

**Figure 2:**
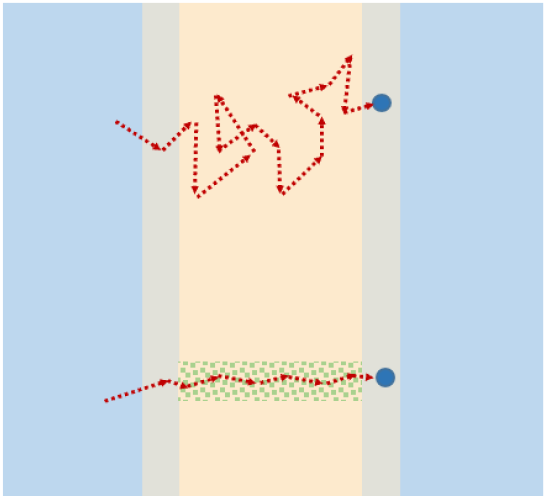
Schematic of the passage of a water molecule by diffusion through the bilayer (top) or through a water channel (bottom).

There is no convective (Navier-Stokes) water flow in the strict sense through aquaporin channels, since the water molecules move in single file. Nonetheless, the molecules are thought to be in close proximity in the channel, and the observed values greater than unity of the 𝒫_*f*_ /𝒫_*d*_ ratio could reflect their influence on each other during osmotic flow.

#### 1.3 Common misconceptions about osmosis

##### Osmotic pressure is not a bulk property of a solution

The major misconception bedeviling our understanding of osmotic pressure and osmosis, for well over a century, is the attribution of these phenomena to the bulk properties of the solutions bathing the membrane, while ignoring the physical implications of the most obvious property of the membrane itself, namely, its mechanical interaction with the solute making it impermeable to the solute molecules. The most common mistake, which has recurred persistently, is the idea that in a free-standing solution, the solute and solvent exert independent pressures, just like a mixture of ideal gases. Modern thermodynamic and statistical mechanical ideas of liquid solutions have fortunately taken root, and today this erroneous picture is only rarely invoked.

##### A gradients of water concentration is not the primary driver of osmosis

Another common misconception corresponds to the explanation that an osmotic water flux across a membrane is caused by a difference in water concentrations between the two solutions separated by the membrane. Here is a typical statement: “Water spontaneously moves ‘downhill’ across a semi-permeable membrane from a solution of lower solute concentration (relatively high water concentration) to one of higher solute concentration (relatively low water concentration), a process termed osmosis, or osmotic flow. In effect, osmosis is equivalent to ‘diffusion’ of water across a semipermeable membrane.” (Lodish et al., 2021). Or, “If cells are placed in a hypotonic solution (that is, a solution having a low solute concentration and therefore a high water concentration), there is a net movement of water into the cells.” (Alberts et al., 2015).

The idea that in osmosis water flows down its concentration gradient is misguided. The difference in water concentration between pure water and an aqueous solution is not simply a function of solute concentration alone. A straightforward calculation shows that it also depends on the ratio of the partial molar volume of the solute species to the molar volume of pure water (see Appendix 3). This ratio is specific to the particular solute species. The same concentration of solute, but for different solute species, leads to water concentration differences between the two solutions that are specific to the specific solute species. If the osmotic flow were caused by the difference in water concentrations, the water flux would then be specific to the solute species used to establish it. Such a dependence on impermeable solute species is not observed for dilute solutions, and moreover, would contradict both van’t Hoff’s law and the Fundamental Law of Osmosis.

##### Osmotic transport is not different from transport induced by pressure differences

The other misconception is to deny the reality of the pressure underlying the movement of water across a semipermeable membrane. Here is an example: “The relationship (van’t Hoff) however arises directly from the parallels in the thermodynamic relationships and should not be interpreted in the molecular mechanistic sense, since the osmotic pressure is in fact a property ensuring equilibrium of the *solvent* and solute, and has its effect only via its reduction of the chemical potential of this solvent.” (Tombs and Peacocke, 1974). The identification of hydrostatic pressure driven flow and flow driven by a concentration imbalance of impermeable solute is embodied in the Fundamental Law of Osmosis, and we will demonstrate how the Debye model explains this equivalence in a physically transparent way.

Erroneous notions of the cause of osmotic flow put about in textbooks, have the unfortunate effect of actively spreading confusion. If progress is to be made in comprehending osmotically driven water fluxes, it is important to ensure that our understanding is firmly rooted in well-established physical principles.

### 2 A mechanistic model for osmosis: the Debye model

Several different mechanisms have been proposed to explain how osmosis arises, with Guell (1991) listing fourteen.^5^ We will argue that there is in fact only one parsimonious explanation for osmosis, that relies on the mechanical interaction between the membrane and impermeable solute molecules, and that we will refer to as the **Debye model**, as it was first proposed by Debye in 1923. Despite Debye’s reputation, the model made little impact on our understanding of osmosis – disappearing for decades, probably because biologists were not aware of it, and chemists and physicists largely uninterested – until the 1960s. Unfortunately the connection to the original work was lost, and we re-establish it here (see the box for a short history).

Debye recognized that the physical principles underlying the development of an osmotic pressure must be centered on the interactions of the membrane with the solute molecules, since osmotic pressure is not observed in the absence of a membrane. As Debye put it in his 1923 paper, “We express the quality of semi-permeability of the membrane by saying that the potential energy of a solute molecule increases from zero to infinity when it is transported across the membrane from the solution side.” An equivalent statement would be that the membrane exerts a repulsive force *ℱ* on a solute molecule that is strong enough to prevent the solute molecule from entering the membrane and crossing over to the pure solvent side.

#### A short history of the Debye model

The investigation of osmosis has an interesting history that has been told by others (Hammel and Scholander, 1976; Mason, 1991). In this section we will focus on the history of the Debye model.

Although the experimental demonstration of osmosis (1748) predates that of diffusion (1828), the development of the theoretical basis of diffusion proceeded with little controversy (Einstein, 1905; Jacobs, 1935; Berg, 1993). In contrast, the theoretical underpinnings of osmotic pressure proved contentious from the start. ^6^

There is a fascinating story recounted by George Wald (1982) that it was Hugo de Vries (a botanist and one of the re-discoverers of Gregor Mendel’s work) who told van ‘t Hoff about Pfeffer’s experiments (Pfeffer, 1890) on semipermeable membranes when their paths crossed while walking in Amsterdam. Van ‘t Hoff was awarded the first Nobel Prize in Chemistry in 1901 largely for his work on osmosis. At our historical remove, it may seem strange to award the prize for what seems like such a simple finding. However, it provided one of the first experimental confirmations of atomic theory. What we have called the Debye model was first proposed by Peter JW Debye in a paper first published in French (Debye, 1923b) and then in German (Debye, 1923a), and primarily devoted to further developments of Debye’s theory of ionic solutions. Debye remarks in a footnote “Among the large number of authors who have already dealt with the kinetic theories of osmotic pressure, we must cite above all: L.Boltzmann, H.A.Lorentz, Ph. Kohlstamm, C.Jäger, O. Stern, P. Langevin, J. J. van Laar, P. Ehrenfest.”, but does not cite any of their papers, because they failed to pin down the mechanism.

In the intervening years there have been very few references to this work. Joos developed a simplified derivation of the mechanism in what is essentially a didactic paper (Joos, 1941), acknowledging that his work was derived from an idea in Debye’s 1923 paper. The derivations were included in his influential textbook of physics (Joos and Freeman, 1959). Manning (1968) was probably the first to re-derive the Debye model in the second half of the 20th century. Manning based his derivation on a textbook by Rutgers (1954), who said that his argument derived from Debye, but Rutgers does not quote the paper. It is worth noting that Debye provided a foreword to Rutgers textbook.

The textbook by Villars and Benedek (1974) is the source most often quoted for the solute-membrane repulsion model, but it has no references at all to Debye, and they likely found it in Manning (1968). In the biological literature Mauro (1979) appears to be the first to have referred to Villars and Benedek in the context of osmosis.

It is puzzling that Debye’s work on osmosis made little impact, since he was a major figure in the development of physics in the twentieth century, receiving the Nobel Prize in 1936. It is even more so because he was a professor at Cornell University (Ithaca, NY, 1939-66) during the period when the debate about the molecular origins of osmosis was revived. Indeed, from the mid-1950s to 1990s several theories competed about the origin of osmotic pressure (Hammel, 1979; Hildebrand, 1979; Mauro, 1979; Soodak and Iberall, 1979; Yates, 1979; Essig and Caplan, 1989). Prominent among the contesting theories was the controversial solvent tension theory (Hammel and Scholander, 1976). However, the Debye model never seemed to have made an appearance in the debate, at least in its quantitative form. In an interview in 1964, Debye himself provides a possible key to this enigma. When asked which periods of his work stand out to him “… I think they are important at the moment when I am doing them. Later I forget about them. So it’s only during the time that I have fun with them that they seem important.” (Corson et al., 1964).

#### 2.1 The Debye model leads to van’t Hoff’s law

Debye was concerned only with osmotic equilibrium, so we begin by following his derivation of van ‘t Hoff’s law for osmotic pressure at equilibrium. Afterwards, we discuss steady state osmotic flow, as a straightforward extension of his model (Manning, 1968). Consider a semipermeable membrane separating two chambers at equilibrium, with the *x* coordinate increasing from left to right, the semi-permeable membrane perpendicular to the *x* axis, the solution compartment with solute concentration *c*_*s,r*_ to the right of the membrane, the pure solvent to the left. We are in effect looking at an infinite 2D membrane, with all values isotropic in the *y* and *z* directions.

Our first goal is to obtain an equation characterizing the solute concentration profile *c*_*s*_ (*x*). For that, we write an equation for the flux *j*_*s*_ of solute molecules in the absence of an applied pressure,

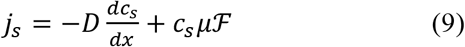

where the first term on the right hand side of the equation is Fick’s law for solute flux in the presence of a solute concentration gradient in dilute conditions, and the second term is the contribution to the solute flux from the mechanical force *ℱ* exerted by the membrane on nearby solute molecules. Einstein’s relation *D* = *RTµ* (Einstein, 1905) will allow us to convert the solute mobility *µ* to its diffusion coefficient *D*. The semi-permeability property of the membrane means that passage of solute into and through the membrane is completely blocked by the force *ℱ*. Therefore, there must be a gradient of solute concentration across the membrane-solution interface where from left to right the solute concentration increases steeply from zero just inside the membrane to the constant value *c*_*s, r*_ of solute concentration in the solution chamber. Moreover, since the membrane excludes the solute, the solute flux across the interface must vanish. Setting *j*_*s*_ = 0, then using Einstein’s relation and cancelling *D*, we obtain an equation to characterize the solute concentration profile *c*_*s*_ (*x*),

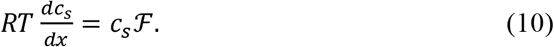

To connect this equation to the pressure that develops across the membrane, we can visualize a volume element of solution near the membrane as a thin slice of thickness *dx* parallel to the membrane (see rectangular blue box in Fig. 3). When the system is at equilibrium, the slice in particular must be in mechanical equilibrium, meaning that all of the forces acting on and inside the slice must balance out to zero. The intermolecular forces among the molecules inside the slice cancel each other as a consequence of Newton’s law of action-reaction, leaving the requirement that the forces on the slice originating from outside it must balance to zero. These forces are the repulsive force *ℱ* from the membrane acting on each solute molecule in the slice, and the hydrostatic pressures from the fluid surrounding the slice and pushing from outside the slice on each of the side surfaces of the slice. With *PP*(*x*) the pressure at *x*, the zero balance is expressed by the equation, *dP*/*dx* − *c*_*s*_ *ℱ* = 0, or,^7^

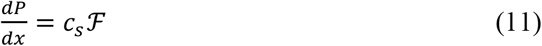

**Figure 3:**
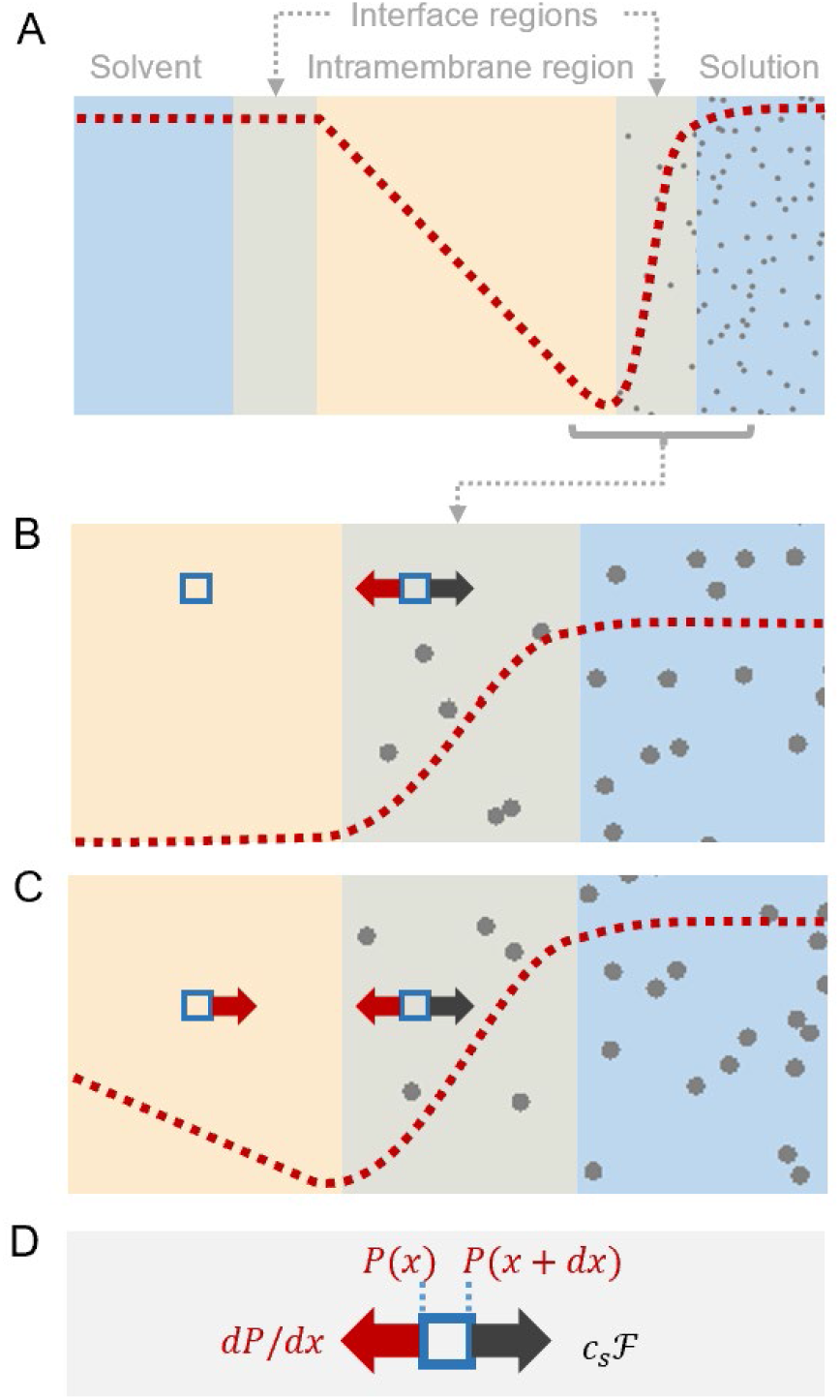
An illustration of the Debye model and the Vegard pressure profile. (A) Schematic view of the cross-section of a membrane illustrating the expected pressure profile (red), with solute molecules (gray) on the right, for the osmotic steady state. (B) and (C) represent magnified views of the solute side of the membrane-solution interface. The blue squares depict a volume element of the solution, with the expected forces shown for the case of the (B) osmotic equilibrium and (C) the osmotic steady state. (D) Shows the forces operating on the volume elements.

Eqs 10 and 11 can be combined,

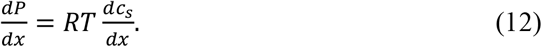

The van ‘t Hoff law for osmotic equilibrium

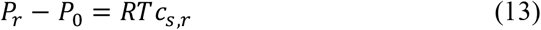

follows after integration from left to right (pure solvent to solution) with *P*_*r*_ the pressure in the solution compartment, *P*_0_ the pressure in the pure solvent compartment, and of course *c*_*s,l*_ = 0 in the pure solvent compartment.

We are now in a position to recognize the genius of Debye’s insight, simple as it is. At the heart of his derivation is the membrane-solute force *ℱ*, which would be different for every membrane and every solute. How can it lead to van’t Hoff’s law, which is applicable generally to any membrane-solute pair? The reason, as we have just seen, is that it produces compensating physical effects and cancels from the final result.

#### 2.2 The Vegard pressure profile

We now move from considering osmotic equilibrium, to the situation where the pressures in the chambers are constrained to be the same, and both chambers are very large and well stirred. Under these conditions, which we will refer to as the **osmotic steady state**, an osmolarity gradient across the semipermeable membrane will drive a steady flow of water across the membrane. We will show that extension of the Debye model to osmosis demonstrates that there must be a pressure drop from the solution to just inside the membrane equal to *RTc*_*s*_. Since the pressure is lower on the solution side (just inside the membrane) than on the pure solvent side, there is necessarily a pressure gradient across the membrane. In a simple one-dimensional visualization the expected pressure profile is shown in Fig. 3 However, the pressure gradient within the membrane may have a more complicated form shaped by the molecular structure of the membrane.

In a prescient 1908 paper, Lars Vegard, who was a student of JJ Thompson at the time, appears to have been the first to propose this pressure profile (Vegard, 1908). He suggested, based on osmotic transport measurements with synthetic membranes, that somehow the solute generated a pressure gradient within the membrane but did not hazard a mechanism. Such pressure profiles were rediscovered by several workers (Dainty, 1965; Mauro, 1965), and in 1968 (Manning, 1968) Manning first made the connection between the profile and the Debye model (see Fig. 3c of (Manning, 1968)). We term this peculiar pressure profile the **Vegard pressure profile**, and the pressure drop in the narrow interface region on the solution side the Vegard pressure drop.

The Vegard pressure profile provides a graphic description of the mechanism that drives the osmotic flow of water. The intramembrane pressure gradient drives water from the side with the lower osmolarity (pure solvent in Fig. 3) to the side with the higher osmolarity. In the narrow interface region on the high osmolarity side, the pressure drop by itself would drive water back towards the membrane, but in this region it is balanced by the forces from the membrane that drive the impermeable solute molecules away.

#### 2.3 The Vegard pressure drop drives osmosis

With reference to Eq. 11 and the discussion above it, we have explained that the difference *dP*/*dx* − *c*_*s*_ *ℱ* represents the net force on a volume element of solution near the membrane-solution interface, and that at equilibrium it equals zero. When the system is not in equilibrium, the difference 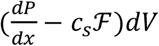 is still the total net force on a volume element *dV* at the membrane-solution interface, but it is not zero and gives rise to a volume flux Φ_*V*_. If the flux is not too large, we can set down a linear relationship between the net force and the volume flux,

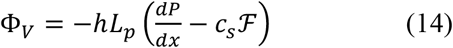

where we will verify the identification of the proportionality constant as *hL*_*p*_, where *h* is the width of the membrane, and *L*_*p*_ is the permeability in Darcy’s Law, Eq. 2. The relation between the membrane force *ℱ* and the solute concentration gradient at the membrane-solution interface, Eq. 10, remains valid in the steady state case, since we assume well-stirred conditions at the interface, so that this expression for Φ_*V*_ becomes,

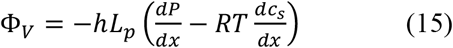

Now we integrate both sides of this equation across the membrane-solution interface from just inside to just outside. The integral involving the volume flux Φ_*V*_ is small, because it is proportional to the narrow width of the interface. But the integrals of the pressure and concentration derivatives do not depend on the width of the interface. The integral of the pressure derivative across the interface equals the pressure difference across the interface. The integral of the derivative of solute concentration equals the difference of solute concentrations across the interface. This latter difference equals the bulk solute concentration in the solution, because the concentration of impermeable solute just inside the membrane is zero. The result then of integrating both sides of Eq. 15 across the membrane-solution interface is that from outside to inside there is a pressure drop equal to *RTc*_*s, r*_ at the interface. In other words, the pressure just inside the membrane on the solution side is lower by this amount than the pressure *P*_0_ of the solution outside. Since the pressure is *P*_0_ in both chambers outside the membrane, there must be a pressure gradient across the entire membrane from *P*_0_ on the pure solvent side to *P*_0_ − *RTc*_*s, r*_ on the solution side, and hence we have produced the Vegard pressure profile and pressure drop.

We can take the derivation one step further, and in doing so, both illuminate the action of the pressure gradient and verify the choice of coefficient *hL*_*p*_. The solute concentration is zero inside the membrane, and so its gradient is also zero there. Setting 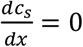 in eq 15, we see that inside the membrane,

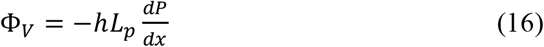

an equation that explicitly exhibits the volume flux as driven by a pressure gradient inside the membrane when the pressures in both solution and pure solvent compartments are equal. Moreover, with these coefficients, this equation is equivalent to Darcy’s law (2).

### 3 Applications of the Debye Model

#### 3.1 The Finkelstein “oil” membrane

As might be expected from van’t Hoff’s law itself, the Debye model does not require detailed knowledge of how water interacts with the specific features of the membrane interior. For example, it is applicable to membranes allowing the passage of water only in the form of isolated independent molecules. To illustrate the Debye model in this context, it is useful to consider a hypothetical scenario that was proposed by Finkelstein (1987). He envisioned a thin “oil” membrane separating two chambers of water. Water is sparingly soluble in the membrane while the oil molecules can diffuse within the plane of the membrane allowing water molecules to move through. We assume that the membrane is stable and cannot be deformed, which is not entirely realistic, but acceptable for theoretical explorations. We show in Appendix 4 that the Vegard pressure drop for this membrane leads to the result *P*_*f*_ = *KD*/*h*, where *K* is the partition coefficient (the ratio of water concentration inside the membrane to that outside), *D* is the diffusion constant of independent water molecules inside the membrane, and *h* is the thickness of the membrane. Since it is clear from inspection that the same result holds for *P*_*d*_, we conclude that *P*_*f*_ /*P*_*d*_ = 1. Since *P*_*f*_ /*P*_*d*_ = 1 is the experimentally measured value for lipid bilayers, the oil membrane model is a useful approximation for biological membranes.

#### 3.2 Single file flow

The experimental observation that 𝒫_*f*_/𝒫_*d*_ exceeds unity for water channels in biological membranes, implies that the unconstrained diffusion of water cannot account for its passage through a protein channel. In the latter, water molecules move in single-file through a narrow passage where water molecules cannot slip past one another. The hypothesis that water channels might restrict the passage of water to move in single file (Lea, 1963), predates the discovery of aquaporin (Preston et al., 1992). This prediction was confirmed by the crystal structure of aquaporin (Walz et al., 1997). However, the precise form of the theory that best accounts for the dynamics of single-file transport of water, in our estimation, remains unresolved (Manning, 1976; Finkelstein, 1987; Horner and Pohl, 2018).

It is worth emphasizing that the Debye model accounts for the osmotic flow through aquaporins, just as it does through any semipermeable membrane. There are Debye/Vegard pressure drops at both ends of the channel, with the larger drop occurring at the end abutting the solution of greater osmolarity. The two ends of the channel face unequal pressures, and the water molecules in the interior of the channel are therefore subjected to a force directed toward the lower of the two pressures.

## 4 Discussion

Our primary objective in this paper is to provide a compelling argument for the Debye model grounded in well-established principles of physics. It begins with the Fundamental Law of Osmosis which implies that whatever happens to drive water across a membrane in the presence of an osmotic gradient, must be the same as for the pressure driven flow in the absence of an osmotic gradient. The Vegard pressure drop, on the side of the membrane adjacent to the solution with the higher osmolarity, provides a plausible mechanical basis for the Law, since the osmotic flow is then also pressure-driven.

A number of scientists have given verbal accounts that accord well with the Debye mechanism, and are worth recalling: “To the extent that it is possible to visualize molecular events, this process could perhaps be pictured (at least for narrow pores) as a molecular piston pump, with solute molecules playing the role of the piston” (Dainty and Ferrier, 1989). And from the great epithelial physiologist Hans Ussing: “The pore contains pure water all the way through, so the driving force cannot be a difference in the chemical activity. Obviously, the driving force is the ‘suction’ created by the osmotic pressure difference at the dotted line. But suction is only another word for hydrostatic pressure difference.” (Ussing and Andersen, 1955)

Physiologists often refer to what is termed the “**colloid osmotic (or oncotic) pressure**”, which is the osmotic pressure that can be attributed to proteins or other large molecules in blood plasma (Boron and Boulpaep, 2016). As blood flows into a capillary bed the hydrostatic pressure filters plasma into the interstitial fluid leaving behind the impermeant proteins in the blood. This has the effect of decreasing the osmolarity of the interstitial fluid relative to the blood. As blood flows out of the capillaries, the hydrostatic pressure declines, and now the osmotic gradient across the capillary wall drives interstitial fluid back into the blood. This interaction between hydrostatic and osmotic gradients, which is of immense importance in clearing the interstitial space, was first postulated by Starling (The fundamental law of osmosis). Although the term ‘colloid osmotic pressure’ is useful in physiology, its mechanistic origins can also be accounted for by the Debye model.

Our focus has primarily been on the physical basis of osmosis, but there are several allied phenomena and concepts that we have not touched on, which are worth mentioning for readers interested in exploring further ramifications of osmosis, namely; depletion forces (Asakura and Oosawa, 1958), diffusioosmosis (Marbach and Bocquet, 2019), osmotic stress (Parsegian, 2002), reflection coefficients (Finkelstein, 1987), and virial coefficients (Neal et al., 1998).

The history of attempts to find a molecular basis for osmosis is surprisingly long and tangled for what on the surface seems like a simple phenomenon. One of the primary difficulties with establishing the physical basis of osmosis is setting up the initial scenario and isolating the essential forces at play. The picture that emerged from the Debye model raised hackles and unfounded thermodynamic arguments were used to counter it. What made this situation even more complicated is that there appeared to be no way of testing the predictions of the theories. After a flurry of activity with no resolution, the debate died out, leaving the erroneous water concentration gradient model uncontested in some textbooks. An odd element that added to the confusion is that even wrong arguments led to the van’t Hoff equation.

It is worthwhile comparing the evolution of our understanding of diffusion to that of osmosis. In the case of diffusion, Einstein’s explanation in 1905 was rapidly confirmed by Jean Perrin’s experiments in 1909 (Perrin, 1909; Perrin, 1910). In contrast, it has taken a very long time for a consistent mechanistic account of osmosis to emerge. To add to that, the scarcity of experiments, and the absence of experiments resolving osmotic mechanisms at the nanometer scale, have perhaps retarded the acceptance of the Debye model.

Molecular dynamics provides a method for exploring what occurs at a molecular level in a phenomenon like osmosis (Roux, 2021). In molecular dynamics, which is now a well-established discipline in molecular physics, Newton’s laws of motion are used to computationally model the collisions of individual molecules. Molecular dynamic simulations using simple particles to represent the solvent and solute together with an energy barrier to model the membrane, successfully recapture van’t Hoff’s law (Murad and Powles, 1993; Zhu et al., 2002; Luo and Roux, 2010; Lion and Allen, 2012). This confirms that the nature of the solvent and solute are irrelevant in generating an osmotic flux. However, molecular dynamics has not been used to model the Vegard pressure profile in steady state osmosis, but it has been used to predict 𝒫_*f*_ / 𝒫_*d*_ from the molecular structure of aquaporins (Zhu et al., 2002; Portella and De Groot, 2009).

With the development of techniques that allow one to probe below the nanometer scale, the precise molecular mechanics of osmotic transport and the validity of the Debye model should be within reach of experiments. It is not inconceivable that molecular sensors could be designed to detect the pressure gradient’s presence and extent. It should therefore be possible to probe the pressure profile first postulated by Vegard in 1908, to confirm a simple and unified view of the physical basis of osmosis.

## Appendix

## Appendix 1: The Donnan Effect

Since the Donnan effect plays an important role in water transport in cells it is worthwhile delving into its nature. To do this we consider a simplified model introduced by Post and Jolly (1957). Let us consider a spherical ‘cell’ with a pliant membrane that is permeable to an uncharged molecule *A* and water. If we place the cell in an infinite bath with *A* at a concentration of [*A*]_*e*_, and assume that the passage of *A* into and out of the cell is governed by the same first-order rate constant *k*, then:

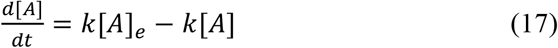

Therefore at equilibrium:

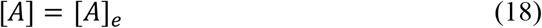

Since the osmolarities inside and outside the cell are balanced, at equilibrium the water flux will be zero and hence the cell will be stable. Now if we introduce *b* moles of an uncharged impermeant molecule *B* into the cell, equation 17 remains unchanged, but the equation for osmotic equilibrium becomes:

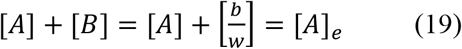

Where *w* is the volume of the cell. The cell must follow the osmotic constraint and the kinetic constraint, and the only way that it can do this is if *w* → ∞. So, water flows in continuously and the cell volume grows without cease. Although we have shown the case for an uncharged molecule, the same holds true for charged solutes. The volume can be stabilized by introducing an impermeant molecule in the extracellular space. However, this is not what animal cells do, instead they pump molecules out of the cell to stabilize cellular volume(Tosteson and Hoffman, 1960). In the case of the toy model that we have introduced here, it can be shown that if *A* is pumped out of the cell, the volume can be stabilized in the presence of *B*.

## Appendix 2: Identification of 𝒫_*d*_

Eq. 7 is a statement of Fick’s law for the diffusion of tracer molecules when there is a concentration gradient of tracer. Therefore 𝒫_*d*_ = *D*/*h*, where *D* is the self-diffusion constant of water molecules in the membrane, and *h* is the membrane thickness. The question is whether this 𝒫_*d*_ also characterizes the diffusive component *ϕ*_*w,d*_ of water flux, not tracer molecules, when a force per molecule *f* is imposed. The answer is yes, as we show here. Note that an arbitrary multiplicative factor, such as a partition coefficient, does not affect the conclusion.

Using the Einstein relation between diffusion constant and diffusional mobility, we have *ϕ*_*w,d*_ = (*D*/*RT*)(*N*_*w*_/*Ah*)*f*, where *N*_*w*_ is the number of water molecules in the membrane, and *Ah* is the volume occupied by the membrane, *A* being the cross-section area and *h* the membrane thickness. The total force *F* on the water in the membrane is *N*_*w*_*f*, and *D*/*h* = 𝒫_*d*_. Then, *ϕ*_*w,d*_ = 𝒫_*d*_(*F*/*A*)/*RT*), where here we use a molar flux. The net force per unit area can be imposed by a pressure difference, and then *ϕ*_*w,dd*_ = −𝒫_*d*_ Δ*P*/*RT*, completing the proof.

## Appendix 3: Relation between water and solute concentrations

The sum of the water and solute concentrations *c*_*w*_ + *c*_*s*_ is (*n*_*w*_ + *n*_*s*_)/*d*, where *n*_*i*_ is the number of moles of species *ii* and *d* is the volume of solution, equal to (*n*_*w*_ *v*_*w*_ + *n*_*s*_*v*_*s*_), where *vv*_*ii*_ is the partial molar volume of species *i*. A straightforward rearrangement leads to

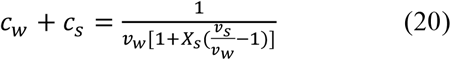

where *X*_*s*_ (= *n*_*s*_ /(*n*_*w*_ + *n*_*s*_)) is the mole fraction of solute. Only if the solute species is essentially identical to water, for example, D _2_O, can we say 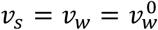, where the latter is the molar volume of pure water, and thus obtain from this equation the simple result *c*_*w*_ + *c*_*s*_ = 55.5*M*.

In this situation the water concentration depends only on the solute concentration and is independent of the specific solute species. In general however, the concentration of water and that of solute are not simply related.

## Appendix 4: The “oil” membrane

Consider a membrane that allows water to move only as independent molecules. To start, the membrane is bathed on both sides by chambers of pure liquid water at the same pressure. The uniform equilibrated concentration of water molecules inside the membrane is denoted by *c*_*w,m*_. Now let a pressure difference Δ*P* be imposed between the two chambers, so that there is a pressure gradient *dP*/*dx* across the membrane. The force on a water molecule inside the membrane is −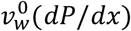. Here we have made a simplifying assumption in the spirit of a mechanical theory. Instead of using a thermodynamically rigorous partial molar volume for water, we have assumed that each water molecule possesses a definite volume, and that this volume inside the membrane is equal to the molecular volume 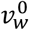 of pure water (the volume of some portion of bulk liquid water divided by the number of water molecules in it).^8^ Using the Einstein relation between the mobility coefficient and the diffusion constant *DD* of water molecules in the membrane, the water flux is − 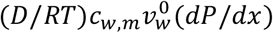. Multiply and divide this expression by 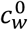, the concentration of pure bulk water (*i*.*e*., 55.5 M), and notice that 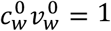. Finally, the molar water flux *ϕ*_*w*_ is obtained on integration across the membrane of thickness *h*,

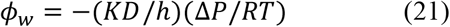

where *K* is the partition coefficient, that is, the ratio 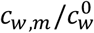 of water concentrations inside and outside the membrane. The sign indicates that the water flows from high to low pressure.

Next, we consider osmotic water flow. The chamber on one side of the membrane is a dilute aqueous solution with solute concentration *c*_*s*_, the solute molecules being impermeable to the membrane. The chamber on the other side is pure water. The water inside the membrane exists as before as independent molecules. There is no pressure difference between the chambers with common pressure *P*_0_, but there is a Vegard pressure drop at the membrane-solution interface equal to *RTc*_*s*_. Therefore there is a pressure gradient in the membrane from low pressure *P*_0_ − *RTc*_*s*_ at the membrane-solution interface, to high pressure *P*_0_ at the interface between membrane and pure water chamber. The water flux must then be identical to the pressure-driven flux with Δ*P* replaced by *RTc*_*s*_,

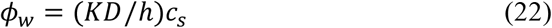

The sign shows that the water flow is into the solution.

In agreement with the Fundamental Law of Osmosis, ((5)), the permeability coefficient 𝒫_*f*_ is seen to be the same either from Darcy’s law ((21)) or from the osmotic flux equation ((22)),

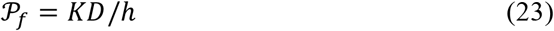

Moreover, it is clear by inspection that the diffusive permeability 𝒫_*d*_ must also be equal to *KD*/*h*, so that 𝒫_*f*_/𝒫_*d*_ = 1.

## Acknowledgements

We thank Sophie Marbach and Hiroaki Yoshida for useful discussions,and Michael J Welsh for helpful suggestions. A.R.K. is supported by NSF grant 2037828.

## Author Contributions

Both authors contributed to the writing of the manuscript and approved its final version. G.S.M. derived the equations, and A.R.K. produced the figures.

The authors declare no competing financial interest.

## Footnotes

1 Van ‘t Hoff’s law Eq. 1 holds for sufficiently dilute solutions. There are different ways of indicating effects caused by interactions among solute molecules at higher concentrations. For simplicity, we discuss dilute solutions only.

2 Darcy’s law, however, has a more general validity. As follows from the very notion of pressure-volume work, application of a Δ*PP* across a membrane that is permeable to water while supporting the pressure must give rise to a transfer of water volume regardless of the physical nature of the water flow inside the membrane.

3 *µ* = 1/*ζ*, where *ζ* is the friction coefficient of the molecules. For a spherical particle with radius *r*, in a solution with viscosity *η*, the Stokes equation holds true *ζ* = 6*πηr*.

4 The measurement of 𝒫_*d*_ is technically challenging, because of the effect of unstirred (unconvected) layers in the juxta-membrane space, which distort measurement of the permeability coefficients. However, it is possible to estimate the size of the unstirred layers and to correct for their influence (Barry and Diamond, 1984).

5 Guell’s list -1) solvent diffusion, 2) solute-membrane collisions, 3) solute suction forces, 4) pore mouth vibration, 5) vaporization and condensation in the membrane pores, 6) solute solvent forces, 7) solute adsorption to the membrane, 8) enhanced solvent tension, 9) reduced solvent activity, 10) free surface solute pressure, 11) solute dissolution in the membrane, 12) membrane steric forces, 13) diffusion pressure, and 14) reflection zones.

6 “Again we have the basically pointless question: What exerts osmotic pressure? Really, as already emphasized, I am concerned in the end only with its magnitude; since it has proved to be equal to the gas pressure one tends to think that it comes about by a similar mechanism as with gases. Let he, however, who is led down the false path by this rather quit worrying about the mechanism. “ - Van’t Hoff (1892) translation from Weiss (1996)

7 If this equation is multiplied on both sides by the volume *Adx* of the slice, with *A* the surface area of the slice, and if the units are then checked while remembering that pressure is force per unit area, it becomes a force balance even more transparently.

8 A more elaborate calculation that does not use this simplification yields exactly the same result.

## References

Agre, P., D. Brown, and S. Nielsen. 1995. Aquaporin water channels: Unanswered questions and unresolved controversies. Curr. Opin. Cell. Biol. 7:472–483.

Alberts, B., A. Johnson, J. Lewis, D. Morgan, M. Raff, K. Roberts, and P. Walter. 2015. Molecular Biology of the Cell. 6th.ed. Garland Press. 1464 pp.

Andersen, O.S. 2015. Perspectives on: The response to osmotic challenges. Journal of General Physiology. 145:371–372.

Asakura, S., and F. Oosawa. 1958. Interaction between particles suspended in solutions of macromolecules. Journal of Polymer Science. 33:183–192.

Baumgarten, C.M., and J.J. Feher. 2011. Osmosis and Regulation of Cell Volume. In Cell Physiology Sourcebook. N. Sperelakis, editor. Elsevier. 261–301.

Benedek, G.B., and F.M. Villars. 1974. Physics with Illustrative Examples from Medicine and Biology. Vol. 2: Statistical Physics. Addison-Wesley Reading, MA.

Berg, H.C. 1993. Random Walks in Biology. Princeton University Press, Princeton, NJ. 152 pp.

Blaustein, M.P., J.P.Y. Kao, and D.R. Matteson. 2019. Cellular Physiology and Neurophysiology. 3rd.ed. Elsevier/Mosby.

Borg, F.G. 2003. What is osmosis? Explanation and understanding of a physical phenomenon. arXiv preprint physics/0305011.

Boron, W.F., and E.L. Boulpaep. 2016. Medical Physiology: A cellular and molecular approach. 3rd.ed. Elsevier. 1312 pp.

Corson, D.R., E.E. Salpeter, and S.H. Bauer. 1964. Peter JW Debye: An Interview. Science. 145:554–559.

Dainty, J. 1965. Osmotic flow. Symp Soc Exp Biol. 19:75–85.

Dainty, J., and J. Ferrier. 1989. OSMOSIS AT THE MOLECULAR-LEVEL. Studia Biophysica. 133:133–140.

Debye, P. 1923a. Kinetische theorie der gesetze des osmotischen drucks bei starken elektrolyten. Physik. Z. 24:334–338.

Debye, P. 1923b. Théorie Cinétique des Lois de la Pression Osmotique des Électrolytes Forts. Recueil des Travaux Chimiques des Pays-Bas. 42:597–604.

Dick, D.A.T. 1966. Cell water. Butterworths.

Dill, K.A., and S. Bromberg. 2003. Molecular driving forces; statistical thermodynamics in chemistry and biology. Garland Science, New York. 666 pp.

Einstein, A. 1905. Über die von der molekularkinetischen Theorie der Wärme geforderte Bewegung von in ruhenden Flüssigkeiten suspendierten Teilchen. Annalen der physik. 17:549–560.

Essig, A., and S. Caplan. 1989. Water movement: does thermodynamic interpretation distort reality? Am J Physiol-Cell Ph. 256:C694–C698.

Fettiplace, R., and D.A. Haydon. 1980. Water permeability of lipid membranes. Physiol Rev. 60:510–550.

Finkelstein, A. 1987. Water movement through lipid bilayers, pores, and plasma membranes. Theory and Reality. John Wiley and Sons., New York. 228 pp.

Guell, D.C. 1991. The physical mechanism of osmosis and osmotic pressure--a hydrodynamic theory for calculating the osmotic reflection coefficient. Massachusetts Institute of Technology, Boston, MA.

Hammel, H. 1979. Forum on osmosis. I. Osmosis: diminished solvent activity or enhanced solvent tension? American Journal of Physiology-Regulatory, Integrative and Comparative Physiology. 237:R95–R107.

Hammel, H.T., and P.F. Scholander. 1976. Osmosis and tensile solvent. Springer Verlag, New York, NY.

Hevesy, G., E. Hofer, and A. Krogh. 1935. The permeability of the skin of frogs to water as determined by D2O and H2O 1. Skandinavisches Archiv fuer Physiologie. 72:199–214.

Hildebrand, J. 1979. Forum on osmosis. II. A criticism of” solvent tension” in osmosis. American Journal of Physiology-Regulatory, Integrative and Comparative Physiology. 237:R108–R109.

Horner, A., and P. Pohl. 2018. Single-file transport of water through membrane channels. Faraday discussions. 209:9–33.

House, C.R. 1974. Water transport in cells and tissues. E. Arnold.

Jacobs, M.H. 1935. Diffusion processes. In Diffusion Processes. Springer. 1–145.

Joos, G. 1941. Zur unterichtsmassigen darstellung des osmotischen drucks. Z. Physik. Chem. Unterricht. 54:65–66.

Joos, G., and I.M. Freeman. 1959. Theoretical Physics. 3rd.ed. Hafner Publishing Co., New York, NY.

Kay, A.R. 2017. How Cells can Control their Size by Pumping Ions. Frontiers in Cell and Developmental Biology. 5.

Kedem, O., and A. Katchalsky. 1958. Thermodynamic analysis of the permeability of biological membranes to non-electrolytes. Biochimica et Biophysica Acta. 27:229–246.

Kiil, F. 1982. Mechanism of osmosis. Kidney international. 21:303–308.

Kramer, E.M., and D.R. Myers. 2012. Five popular misconceptions about osmosis. American Journal of Physics. 80:694–699.

Lea, E. 1963. Permeation through long narrow pores. Journal of theoretical biology. 5:102–107.

Lion, T.W., and R.J. Allen. 2012. Osmosis in a minimal model system. The Journal of Chemical Physics. 137:244911.

Lodish, H., A. Berk, C.A. Kaiser, M. Krieger, A. Bretscher, H. Ploegh, K.C. Martin, M. Yaffe, and A. Amon. 2021. Molecular Cell Biology. 9th.ed. W.H. Freeman. 1264 pp.

Luo, Y., and B. Roux. 2010. Simulation of Osmotic Pressure in Concentrated Aqueous Salt Solutions. Journal of Physical Chemistry Letters. 1:183–189.

Manning, G.S. 1968. Binary Diffusion and Bulk Flow through a Potential-Energy Profile: A Kinetic Basis for the Thermodynamic Equations of Flow through Membranes. The Journal of Chemical Physics. 49:2668–2675.

Manning, G.S. 1976. Deviation from the Einstein relation of the single-file diffusion coefficient. Biophysical Chemistry. 5:389–394.

Marbach, S., and L. Bocquet. 2019. Osmosis, from molecular insights to large-scale applications. Chemical Society reviews. 48:3102–3144.

Mason, E. 1991. From pig bladders and cracked jars to polysulfones: an historical perspective on membrane transport. Journal of membrane science. 60:125–145.

Mauro, A. 1957. Nature of solvent transfer in osmosis. Science. 126:252–253.

Mauro, A. 1965. Osmotic flow in a rigid porous membrane. Science. 149:867–869.

Mauro, A. 1979. Forum on osmosis. III. Comments on Hammel and Scholander’s solvent tension theory and its application to the phenomenon of osmotic flow. American Journal of Physiology-Regulatory, Integrative and Comparative Physiology. 237:R110–R113.

Murad, S., and J.G. Powles. 1993. A computer simulation of the classic experiment on osmosis and osmotic pressure. The Journal of Chemical Physics. 99:7271–7272.

Neal, B.L., D. Asthagiri, and A.M. Lenhoff. 1998. Molecular Origins of Osmotic Second Virial Coefficients of Proteins. biophs. j. 75:2469–2477.

Nelson, P. 2014. Biological Physics: Energy, Information, Life. W.H. Freeman. 600 pp.

Niklas, K.J., and H.-C. Spatz. 2012. Plant physics. University of Chicago Press.

Oster, G., and C.S. Peskin. 1992. Dynamics of osmotic fluid flow. In Mechanics of Swelling. Springer. 731–742.

Paganelli, C.V., and A. Solomon. 1957. The rate of exchange of tritiated water across the human red cell membrane. The Journal of general physiology. 41:259–277.

Parsegian, V.A. 2002. Protein-water interactions. International review of cytology. 215:1–31.

Perrin, J. 1909. Mouvement brownien et réalité moléculaire. Annales de Chimie et de Physique.

Perrin, J. 1910. Brownian movement and molecular reality. Taylor & Francis

Pfeffer, W. 1890. Osmotische Untersuchungen. Leipzig, 1877.: Zur Kenntniss der Plasmahaut und der Vacuolen. Abh. König. Sächs. Gesell. Wiss., Math.-Phys. Cl. 16:185–344.

Phillips, R., J. Kondev, J. Theriot, and H. Garcia. 2012. Physical biology of the cell. Garland Science.

Portella, G., and B.L. De Groot. 2009. Determinants of water permeability through nanoscopic hydrophilic channels. biophs. j. 96:925–938.

Post, R.L., and P.C. Jolly. 1957. The linkage of sodium, potassium, and ammonium active transport across the human erythrocyte membrane. Biochim Biophys Acta. 25:118–128.

Preston, G.M., T.P. Carroll, W.B. Guggino, and P. Agre. 1992. Appearance of water channels in Xenopus oocytes expressing red cell CHIP28 protein. Science. 256:385–387.

Robbins, E., and A. Mauro. 1960. Experimental study of the independence of diffusion and hydrodynamic permeability coefficients in collodion membranes. The Journal of general physiology. 43:523–532.

Rolfe, D.F., and G.C. Brown. 1997. Cellular energy utilization and molecular origin of standard metabolic rate in mammals. Physiol Rev. 77:731–758.

Roux, B. 2021. Computational Modeling and Simulations of Biomolecular Systems. World Scientific.

Rutgers, A.J. 1954. Physical Chemistry. Interscience Publishers, Inc., New York, NY. 804 pp.

Song, L., M. Heiranian, and M. Elimelech. 2021. True driving force and characteristics of water transport in osmotic membranes. Desalination. 520:115360.

Soodak, H., and A. Iberall. 1979. Forum on osmosis. IV. More on osmosis and diffusion. American Journal of Physiology-Regulatory, Integrative and Comparative Physiology. 237:R114–R122.

Starling, E.H. 1896. On the Absorption of Fluids from the Connective Tissue Spaces. J Physiol. 19:312–326.

Stein, H.J. 1966. Osmotic Theory: An Example of Lag between Research and Teaching. BioScience. 16:97–97.

Tombs, M.P., and A.R. Peacocke. 1974. Osmotic Pressure of Biological Macromolecules. Clarendon Press.

Tosteson, D.C., and J.F. Hoffman. 1960. Regulation of cell volume by active cation transport in high and low potassium sheep red cells. J Gen Physiol. 44:169–194.

Truskey, G.A., F. Yuan, and D.F. Katz. 2009. Transport Phenomena in Biological Systems. 2nd.ed. Pearson. 888 pp.

Ussing, H., and B. Andersen. 1955. The relation between solvent drag and active transport of ions. In Proceedings of the Third International Congress of Biochemistry, Brussels.

van’t Hoff, J.H. 1892. Zur Theorie der Lösungen. Zeitschrift für Physikalische Chemie. 9:477–486.

Vegard, L. 1908. On the free pressure in osmosis. Proc. Camb. Philos. Soc. 15:13–23.

Wald, G. 1982. Origin of the Theory of Solutions. Science. 217:1084–1084.

Walz, T., T. Hirai, K. Murata, J.B. Heymann, K. Mitsuoka, Y. Fujiyoshi, B.L. Smith, P. Agre, and A. Engel. 1997. The three-dimensional structure of aquaporin-1. Nature. 387:624–627.

Weiss, T.F. 1996. Cellular Biophysics: Transport. MIT Press Boston, MA. 693 pp.

White, S.H., G. von Heijne, and D.M. Engelman. 2022. Cell Boundaries: How Membranes and Their Proteins Work. Garland Science.

Yates, F.E. 1979. Introducing a forum on osmosis. American Journal of Physiology-Regulatory, Integrative and Comparative Physiology. 237:R93–R93.

Zhu, F., E. Tajkhorshid, and K. Schulten. 2002. Pressure-induced water transport in membrane channels studied by molecular dynamics. biophs. j. 83:154–160.

